# Impact of fluorescent protein fusions on the bacterial flagellar motor

**DOI:** 10.1101/152595

**Authors:** M Heo, AL Nord, D Chamousset, E van Rijn, HJE Beaumont, F Pedaci

**Affiliations:** Single-molecule Biophysics dept, Centre de Biochimie Stucturale, CNRS UMR5048 UM INSERM U1054, 29 Rue de Navacelles, 34090 Montpellier, France; Department of Bionanoscience, Kavli Institute of Nanoscience, Delft University of Technology, 2628 CJ, Delft, The Netherlands

## Abstract

Fluorescent fusion proteins open a direct and unique window onto protein function. However, they also introduce the risk of perturbation of the function of the native protein. Successful applications of fluorescent fusions therefore rely on a careful assessment and minimization of the side effects. Such insight, however, is still lacking for many applications of fluorescent fusions. This is particularly relevant in the study of the internal dynamics of motor protein complexes, where both the chemical and mechanical reaction coordinates can be affected. Fluorescent proteins fused to the *stator* of the bacterial flagellar motor (BFM) complex have previously been used to successfully unveil the internal subunit dynamics of the motor. Here we report the effects of three different fluorescent proteins fused to the stator, all of which altered BFM behavior. The torque generated by individual stators was reduced while their stoichiometry in the complex remained unaffected. MotB fusions decreased the rotation-direction switching frequency of single motors and induced a novel BFM behavior: a bias-dependent asymmetry in the speed attained in the two rotation directions. All these effects could be mitigated by the insertion of a linker at the fusion point. These findings provide a quantitative account of the effects of fluorescent fusions on BFM dynamics and their alleviation—new insights that advance the use of fluorescent fusions to probe the dynamics of protein complexes.

**Author summary:** Much of what is known about the biology of proteins was discovered by fusing them to fluorescent proteins that allow detection of their location. But the label comes at a cost: the presence of the tag can alter the behavior of the protein of interest in unforeseen, yet biologically relevant ways. These side effects limit the depth to which fluorescent proteins can be used to probe protein function. One of the systems that has been successfully studied with fluorescent fusions for which these effects have not been addressed are dynamic protein complexes that carry out mechanical work. We examined how fluorescent proteins fused to a component of the bacterial flagellar motor complex impacts its function. Our findings show that the fusion proteins altered biologically relevant dynamical properties of the motor, including induction of a novel mechanical behavior, and demonstrate an approach to alleviate this. These results advance our ability to dissect the bacterial flagellar motor, and the internal dynamics of protein complexes in general, with fluorescent fusion proteins while causing minimal perturbation.

## 1 Introduction

Fusion of a fluorescent protein to a protein of interest is an invaluable tool for the study of protein function in a broad, and still expanding range of *in vivo* and *in vitro* systems. However, it is widely recognized that the presence of fluorescent proteins can alter functional properties of the native protein [1–5]. Careful assessment of these side effects and strategies to minimize them have therefore been critical to the success of fluorescent protein fusions (FPFs) [6–9]. The continuing development of FPF-based approaches to explore novel phenomena also calls for new insight into the associated side effects and methods to alleviate them [10–12]. One direction of research for which this holds is the study of the internal dynamics of proteins complexes [13,14].

Recently, several studies have made use of FPFs to study the internal subunit dynamics of the bacterial flagellar motor (BFM) [15–29]. Although FPFs were shown to affect BFM function by decreasing the chemotactic motility of cells and average speed of motors [15], a quantitative characterization of the impact on the BFM mechanical behavior is lacking. Such perturbations limit the depth to which the internal dynamics and function of the BFM, and protein complexes in general, can be probed with FPFs. Linker peptides inserted at the fusion point have been instrumental in minimizing non-native behaviour of FPFs [8–11], but their use in the study of protein-complex dynamics is limited for the BFM.

The BFM is located at the base of each flagellum in the membrane of many motile bacteria (Fig 1A). The torque generated by the complex, and the consequent rotation of the flagella, powers chemotactic cellular motility along chemical gradients [30–32]. Chemotaxis is achieved by chemostimulus-controlled modulation of the motor rotational bias. When all motors of the cell spin counterclockwise (CCW), the cell swims in a straight ‘run’, whereas the switch of one or more motors to the clockwise (CW) direction causes the cell to ‘tumble’ [33]. The CCW to CW bias of a single motor is regulated by the intracellular concentration of the response regulator of the chemotaxis-signaling pathway, CheY-P [34–36]. Torque is generated by up to a dozen stator units, which use the cellular ion motive force (IMF) to perform work on the rotor part of the BFM [31]. In the bacterium *Escherichia coli*, each stator is an ion channel composed of four MotA subunits and two MotB subunits. The stators dynamically turn over between two populations: one that is bound to the BFM and one that passively diffuses in the inner membrane [15,37]. These dynamics have been observed directly at the single-stator level using fluorescent fusions to the N-terminus of MotB [15,18,19, 21, 28, 29]. More recently, similar fusions have been used to unveil that the number of stators recruited into the complex depend on the load of the BFM [18,19, 29].

**Fig 1.**
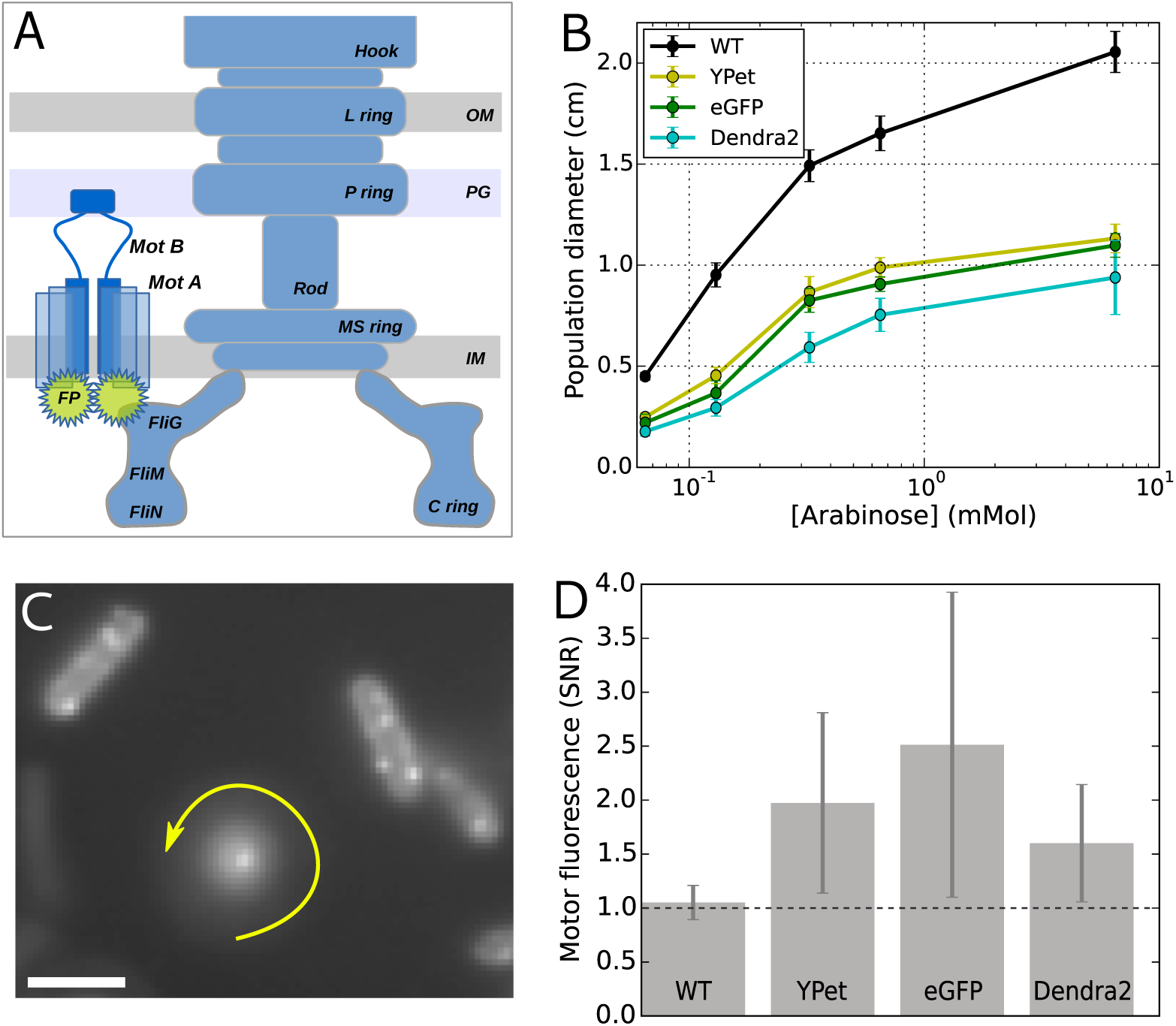
A) Schematic structure of the BFM with one stator labeled by a fluorescent protein fused on the N-terminus of MotB (OM: Outer membrane, PG: peptidoglygan, IM: internal membrane). B) Population chemotactic motility on soft agar plates. The diameter of the ring formed by motile populations after 8h is shown as a function of stator induction, showing a decreased chemotactic motility in strains with labeled stators (error bars indicate standard deviation). C) Stator fluorescence detection from an active labeled motor. In the center, a tethered cell rotates around the bright spot corresponding to the tagged BFM (eGFP-MotB here). The image is the sum of several frames, therefore blurring indicates the cell rotation. In each frame, the rotating cell is visible with different orientations. Fluorescently tagged motors are also visible as bright spots in cells stuck on the glass surface. Scale bar: 2 *μ*m. D) Signal to noise ratio (SNR) for motor fluorescence detected exclusively in rotating tethered cells as in C ([Arabinose]=0.13 mM). Absence of fluorescence is reflected by the line SNR=1 (see Fig S1 for further information). Error bars indicate standard deviation.

In large protein complexes such as the BFM and, for example, polymerases or ribosomes, function emerges from a myriad of dynamical interactions between the subunits. While FPFs have proven to be a very powerful tool to study these complexes, it is likely that they impart biologically relevant functional perturbations. Here, aiming to address this, we fused different fluorescent proteins to the N-terminus of MotB and characterized the effects at the level of individual BFMs. The results show that the observed decrease in population-level chemotactic motility is underpinned by changes, some of which unexpected, in key mechanical parameters of the BFM rotation. On the other hand, stator stoichiometry results unaffected by the presence of the label. Furthermore, we show that both choosing the right fluorescent protein and introducing a rigid linker at the fusion point can mitigate all of these side-effects.

## 2 Results

### 2.1 Functional stators, but reduced chemotactic motility

After fusing the fluorescent proteins YPet, eGFP and Dendra2 directly to the N-terminus of MotB (FPs-MotB), as done in several studies with (e)GFP [15,18,19, 21, 29, 38], we verified that the constructed strains were motile in soft agar at the population level (Fig 1B). Previous studies using eGFP-MotB used a construct containing 500bp upstream of and including the first 28 codons of *motB* (encompassing the putative membrane-targeting sequence), followed by *eGFP* and then the first 500bp of *motB* [15,18, 21]. We compared this construct to one in which eGFP was fused directly to the N-terminus of MotB and found that the latter performed similarly or slightly better in tests of chemotactic population motility.

At the two induction levels tested, the chemotactic population motility of cells expressing FPs-MotB was reduced by ~ 50% relative to to cells expressing wild-type MotB, in line with what was previously found for a GFP-MotB fusion [15, 38]. When observed in TIRF, tethered cells were often found to rotate around a fixed bright spot, corresponding to the location of the rotating BFM (one example is shown in Fig 1C). Image analysis of the fluorescence signal of functional motors, taking into account the auto-fluorescence of the cells and the background signal level, shows that the motor signal-to-noise ratio is higher than one in FPs-MotB strains, while remains equal to one in WT (Fig 1D and Fig S1 in the Supporting Information). Together, these results indicate that the motors are functional, that the labeled stators are localized at the motor, and that all fluorescent proteins fused to MotB impacted BFM function.

### 2.2 Symmetric speed reduction and speed asymmetry in tagged motors

A reduction in chemotactic population motility on soft agar for the FP-MotB strains can be the result of different factors, such as decreased BFM torque, a suboptimal switching frequency (tumbling allows cells to escape dead-ends in the agar matrix [39]), a suboptimal response of the BFM to CheY-P concentrations (i.e. the output of the chemotaxis signalling pathway), or changes in other BFM dynamics. To get insight into the effect of the fluorescent tag on the motor, we used a tethered-bead assay to measure the rotational speed of individual motors. Fig 2 shows the speed distributions for single motors, from all the strains, rotating a high load (1.1 *μ*m diameter bead). The speed distributions are colored according to their bias, with blue and red indicating CCW (counter clockwise, positive speed) and CW (clockwise, negative speed) biased motors, respectively. WT (wild type stators, inducible plasmid) and WTNP (wild type stators, native promoter, see Table 1) behave similarly: regardless of their bias, they reach nearly the same absolute value of speed (±50 Hz) in both directions, in line with previous results [40]. In BFMs with wild-type stators, this behavior is not affected by a change in the induction level of MotB (Fig S2).

**Fig 2.**
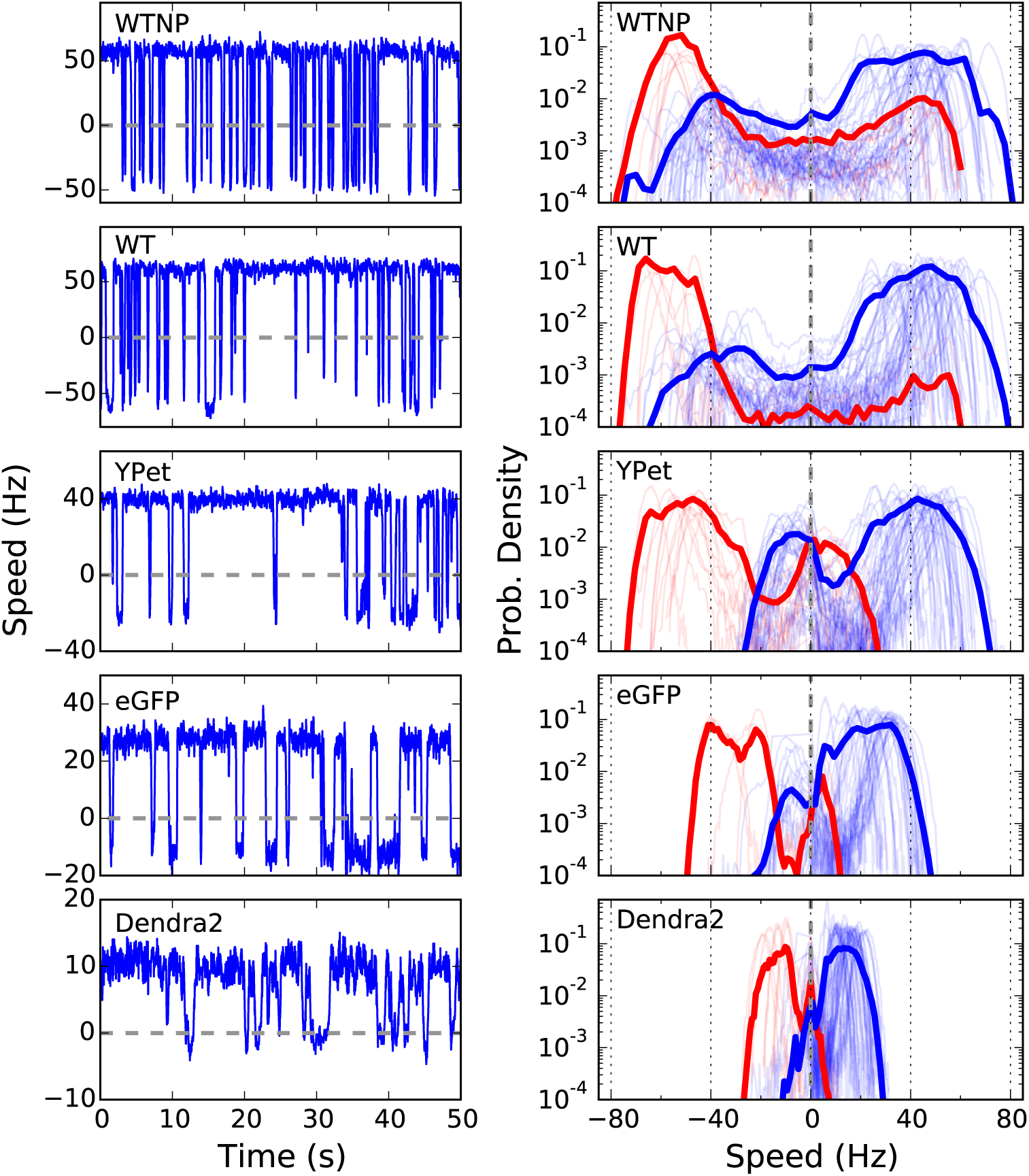
Left column: example speed traces of individual CCW-biased motors rotating 1.1 *μ*m beads. Right column: probability distributions of the speed. Positive and negative speeds indicate CCW and CW rotation direction, respectively. Blue and red indicate CCW and CW biased motors, respectively. Individual thin lines show individual motor measurements, and the thick lines show the average of all measurements. The induction level of stators in WT, YPet, eGFP and Dendra2 is set by an arabinose concentration of 6.5 mM (see Fig S2 for the same measurements at a lower induction level). WTNP: WT with native promoter.

**Table 1.** *E. coli* strains used in this work. All strains come from the parent strain RP437 [53]. FliC^st^ indicates the hydrophobic variant of FliC (producing “sticky” filaments [54]). The fluorescent proteins (FP) fused to MotB are YPet, eGFP, and Dendra2. All plasmids were pBAD33 vector, carrying chloramphenicol resistance and induced by L-arabinose. Strains JPA604, JPA605, and MT02 were gifts from RM Berry lab.

| *Ref.Name* | *Strain* | *Genome* | *Plasmid* | *Assay* |
| --- | --- | --- | --- | --- |
| WTNP | MT02 | *FliC<sup>st</sup>* | – | tethered-bead |
| FP-MotB | JPA605 | *FliC<sup>st</sup>,*<br>*ΔMotAB* | *MotA* FP-*MotB* | tethered-bead,<br>tethered-cell |
| WT | JPA604 | *ΔMotAB* | *MotA MotB* | chemotactic<br>motility |
| FP-MotB | JHC36 [55] | *FliC<sup>st</sup>,*<br>*ΔMotAB,*<br>*ΔCheY* | *MotA MotB* | tethered-bead<br>torque steps |

However, a tagged motor behaves differently. First, regardless of the bias, the most visited speeds of the FP-MotB strains in the two directions are lower than in WT and WTNP. The amount of decrease depends on the tag: YPet reaches almost WT speeds, while eGFP can reach only ~ ±30 Hz, and Dendra2 only ~ ±20 Hz. This *symmetric reduction* of the most visited speed affects motors biased in both directions. Second, a novel feature arises when one considers the bias of the tagged motors. Contrary to WT, when a tagged motor switches from its preferred direction of rotation, it reaches only a fraction of its previous speed in the opposite direction. In FP-MotB strains, this is evidenced in the global average distributions (Fig 2) by the secondary blue and red peaks at negative and positive speeds, respectively. We label this effect *speed asymmetry*. It is observed in a given strain for both biases: tagged CCW-biased motors switch from a high CCW speed to a lower CW speed, while CW-biased motors switch from a high CW speed to a lower CCW speed. The speed asymmetry of tagged motors is therefore bias-dependent, and always results in a lower speed in the less visited direction of rotation.

It has previously been shown that the bias of an individual motor, that is, the percentage of time the motor spends rotating CCW, shows a sigmoidal relationship with the concentration of CheY-phosphate (CheY-P) within the cell [35]. CheY-P is the output of the chemotactic signal transduction network which detects changes in the chemical composition of the environment [41]. This sigmoidal relationship, characterized by a large Hill coefficient, leads to a bimodal distribution of motor bias [42], as seen in Fig 3A. For WT, we measure a switching frequency and distribution of motor bias that is in line with previous results [35, 42]. However, in the tagged motors, we observe a reduction of the frequency of switches with respect to WT (Fig 3A, left panels), with a severity that depends on the particular FP and reflects the order observed above for the decrease in speed (from the least to the most affected: YPet, eGFP, Dendra2). The distribution of motor biases, on the other hand, is less affected (Fig 3A right panels). The reverse cumulative distribution of the residence times (indicating at time *t_o_* the percentage of observed residence times longer than *t_o_* [43]) is shown in Fig 3 for the CCW and CW states. As observed previously, we find that the residence times are distributed non-exponentially with long tails, potentially due to the presence of signaling noise within the chemotaxis network [43]. Reflecting their decreased switching frequency, tagged motors show extended residence times in both directions of rotation with respect to WT.

**Fig 3.**
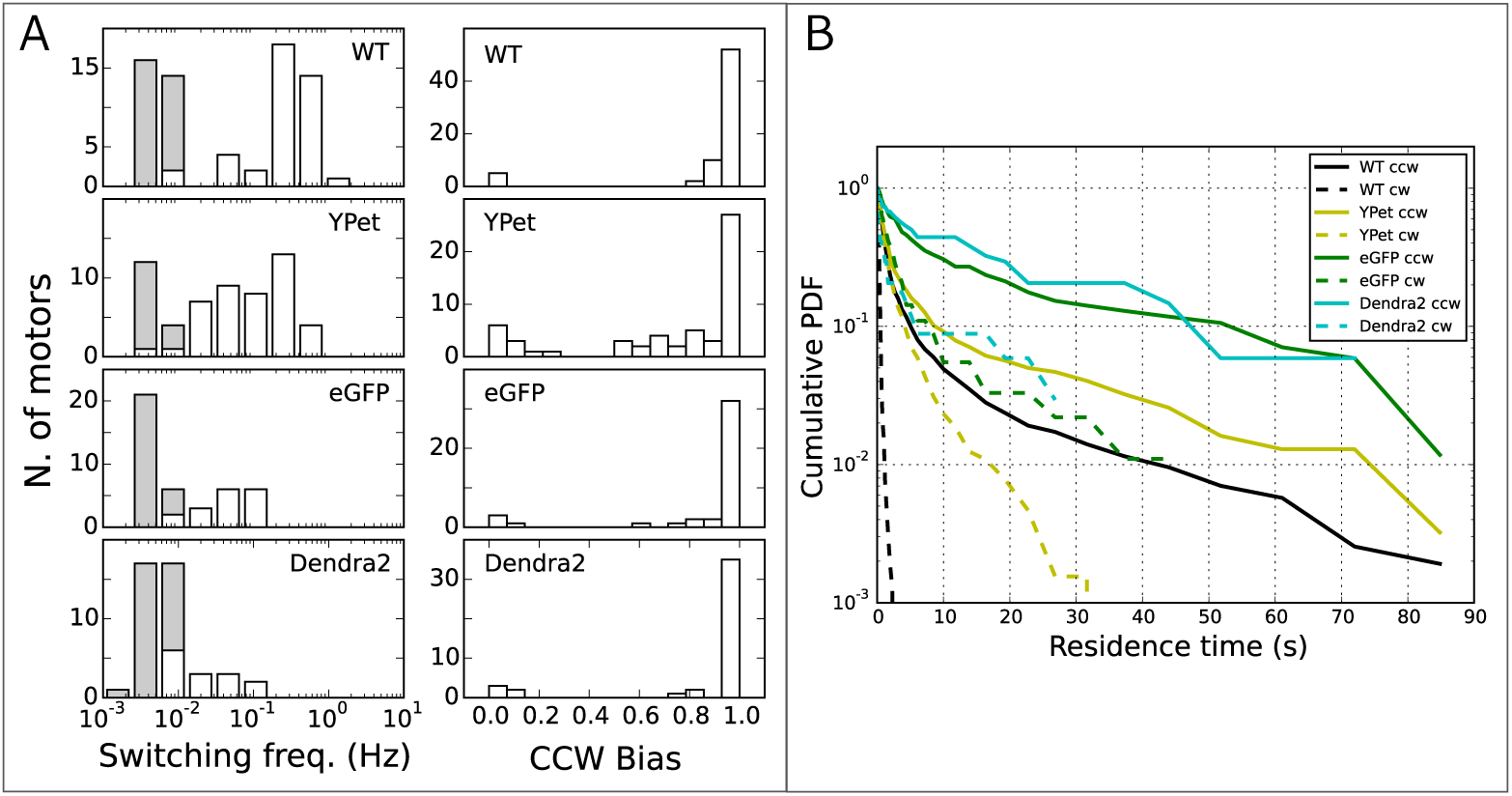
A: Distributions of switching frequency (left column) and CCW bias (right column) for all the strains tested. Gray columns indicate motors which did not switch during the measurement (their switching frequency has been set equal to twice the inverse of the measurement time, and it should be considered as an upper limit). B: Reverse cumulative probability distribution of the time spent in CW and CCW (indicating at time *t_o_* the percentage of observed residence times longer than *t_o_* [43]) for all (CCW biased) motors of the different strains. Stator induction is set by a concentration of arabinose of 6.5 mM (see SI for the same measurements at a lower induction).

### 2.3 Torque and stoichiometry of tagged stators

Decreased populaton-level chemotactic motility and an overall decrease of speed in tagged motors could be due to either a lower torque generated by individual stators or by a lower number of stators bound to the BFM, relative to WT. To discriminate between these two possibilities, we have analyzed the discrete steps in the torque traces which occur spontaneously due to stator turnover [15] in single motors. For this analysis, we used strains lacking the switching regulating protein CheY to avoid the complication of switching events. Fig 4A shows an example torque trace from WT at steady-state, where different torque levels due to stator turnover are detected, and Fig 4B shows the distribution of the torque contributed by a single stator for all the recordings (see Methods and Materials). Gaussian fits to the distributions, reflecting the average torque contribution of a single stator, give the values of 157 ± 43 pN nm in WT, 126 ± 58 pN nm in YPet, 91 ± 30 pN nm in eGFP, and 55 ± 33 pN nm in Dendra2 fusions. Thus, the torque generated by a single stator decreases in the tagged motors, following the same trend observed for the symmetric decrease of speed (Fig 2) and for the decrease in switching frequency (Fig 3).

**Fig 4.**
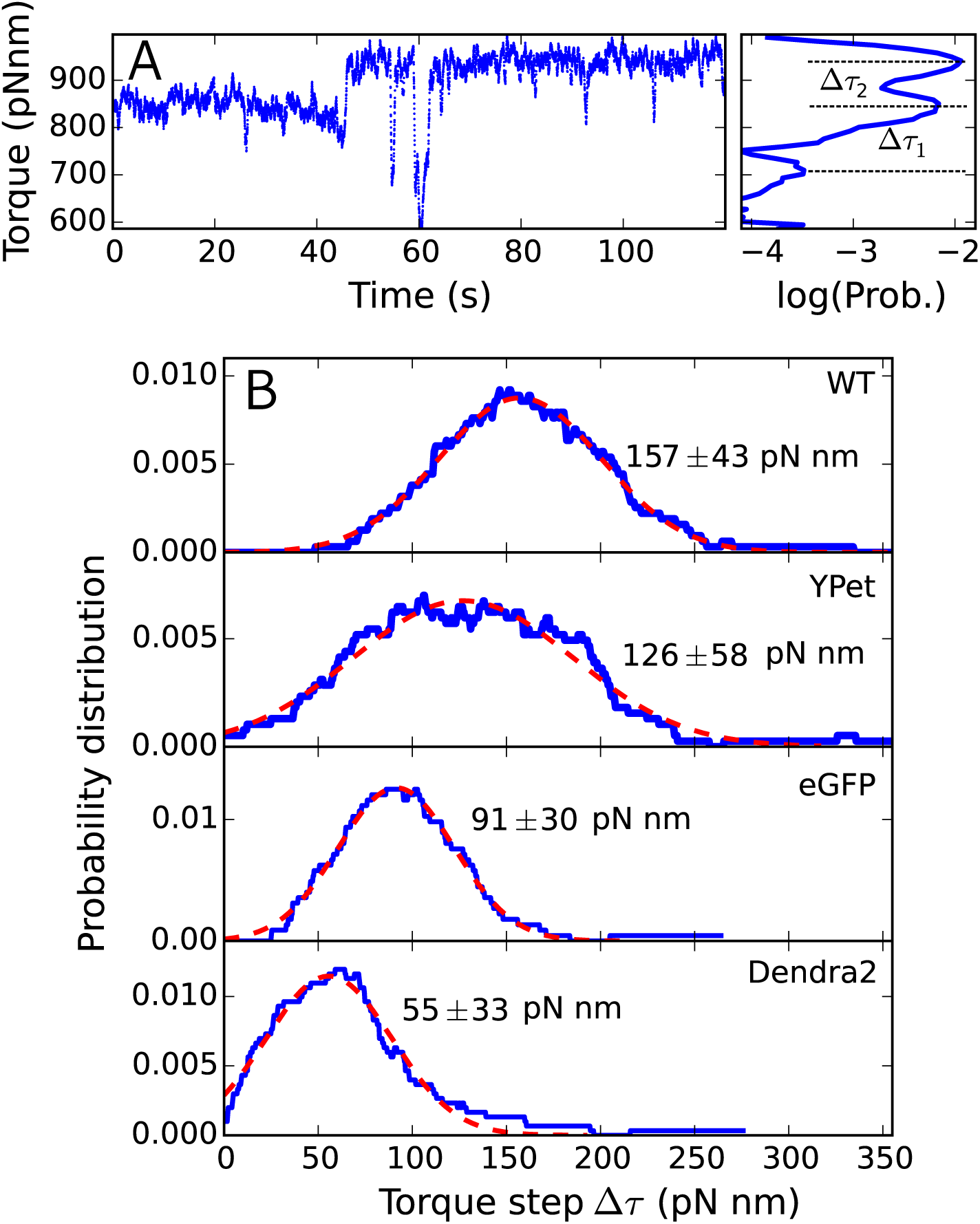
A: An example torque time trace (left) in WT, showing spontaneous stator turnover as steps in torque level. The separation between neighboring torque levels (Δ*τ_i_*) is determined from a multiple Gaussian fit of the torque histogram (right). B: Distributions of the distances between neighboring Gaussians (Δ*τ_i_*) in the multiple Gaussian fit of motor torque, measured in single torque traces like A. The peak indicates the average torque produced by the exchange of a single stator. A single Gaussian fit (red dashed line) provides the center and standard deviation of the peak (indicated by the text in each panel). Stator induction is set by a concentration of arabinose of 6.5 mM.

Moreover, the ratio between the torque per stator in tagged motors and in WT measured here reflects closely the ratio of the most visited speeds (either CW or CCW) in tagged motors and in WT, shown in Fig 2. This implies that the number of stators does not change significantly in tagged motors with respect to WT or WTNP, and that the symmetric speed reduction observed in the tagged motors is mainly the result of the reduced torque generated by each stator. Considering the torque produced by *N* stators as *Nτ*_1_ = *γ*2*πω_N_* (where *τ*_1_ is the torque produced by a single stator, *γ* the drag coefficient of the bead, and *ω_N_* the measured speed (in Hz) of the motor with *N* stators), all the strains tested are driven by *N* ~ 8 – 10 stators, in line with previous measurements at high load [15, 44]. This result supports previous works focused on quantifying steady-state stator stoichiometry by fluorescent stators [15,18,19,21,28,29,45–48].

### 2.4 A linker improves the performance of tagged stators

Aiming to mitigate the effects described above in tagged motors, we introduced a linker between the N-terminus of MotB and the fluorophore. We have tested two types of linkers, one rigid (EAAAK) and one more flexible (GGGGS) [11], both in one copy or as a triple repeat. Population motility measurements, shown in Fig S5, show the results of these tests in Dendra2. Generally, we found that a rigid and longer linker demonstrated a greater improvement in motor performance that a flexible and shorter linker, respectively, with a linker composed of a triple EAAAK repeat showing the greatest improvement.

Focusing on the Dendra2 fusion, which most affects the dynamics of the BFM, in Fig 5 we compare single Dendra2 motors in the presence and absence of the (EAAAK)3 linker (here for CCW biased motors only, due to low statistics for CW bias). In the presence of the linker, the most visited CCW speed increases up to ~ 30 Hz (Fig 5A), while it is only ~ 18 Hz in the absence of the linker (and ~ 50 Hz in WT, see Fig 2). Therefore, the overall speed reduction relative to WT is mitigated by the presence of the linker. We have seen above (Fig 2) that switching in Dendra2 motors is actually more similar to a pause, as the least visited speed remains peaked at zero. In the presence of the linker, motor switching is restored, though a degree of speed asymmetry remains, and the CW rotation is recuperated with a speed peaked at ~ –10 Hz. Fig 5B shows that the higher speed in the presence of the linker is paralleled by a higher torque generated by each stator (with ~ 100 pN nm per stator in the presence of the linker, ~ 55 pN nm in Dendra2 without the linker, and ~ 160 pN nm in WT, see Fig 4). As before, this suggests that the number of stators in the motor is similar in the presence and absence of the linker (*N* ~ 8 – 9), and that the decrease in overall speed is due to a lower torque (and speed) generated by each stator. Finally, Fig 5C shows the change in switching frequency, which increases in the presence of the linker and moves towards that of WT (as shown in Fig 3A). Similar to Dendra2, in YPet and eGFP tagged motors the presence of the linker is responsible for a partial recovery of speed and decrease in severity of the speed asymmetry, as shown in Fig S6.

**Fig 5.**
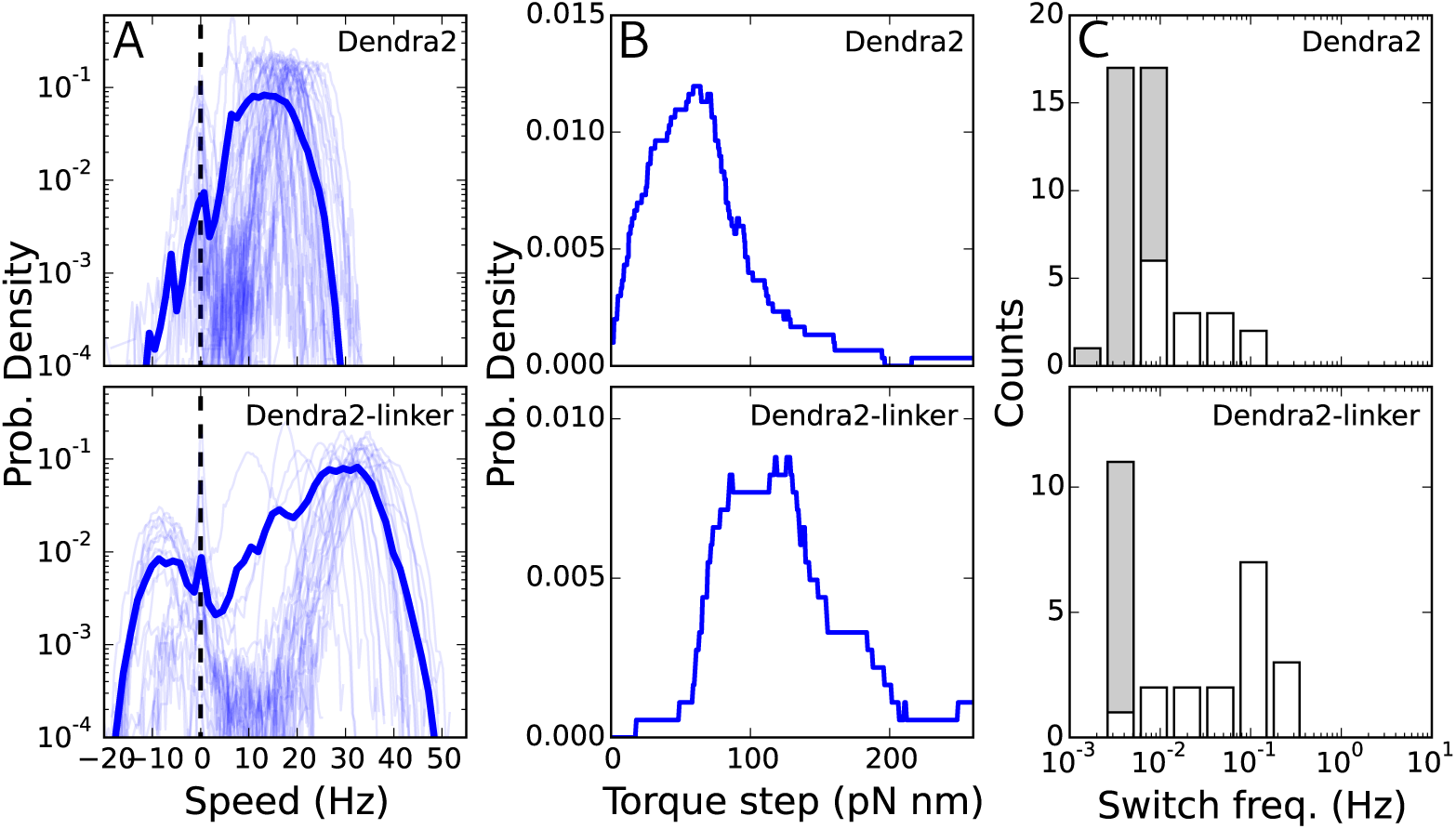
Introducing a linker between the FP and MotB improves the features of the tagged motors. We show here the results for Dendra2-MotB (and in Fig S6 the analysis for YPet-MotB and eGFP-MotB). Top (low) panels correspond to the absence (presence) of the (EAAAK)_3_ linker. A) Probability distribution of the speed (as in Fig 2) of CCW biased Dendra2-MotB motors (CW biased motors are not shown due to low statistics). B) Probability distribution of the torque steps observed at steady-state during stator turnover (as in Fig 4B). C) Distribution of measured switching frequency (as in Fig 3A). [Arabinose] = 6.5 mM.

## 3 Discussion

Fluorescent protein fusions have been successfully used to study the internal dynamics of protein complexes, but insight into the effect of the label on biologically relevant functions of the complex is still limited. We examined this for fluorescent fusions that have previously been used to shed light on the internal dynamics of the BFM [15,18,19, 21, 28, 29]. Globally, we find that the fusion of an FP at the N-terminus of MotB affects several features in the dynamical behaviour of the motor driving a high load. The overall speed of the BFM is reduced due to a reduction of the torque produced by each stator, while the stator occupancy remains unaffected. Also, the switching frequency of tagged motors decreases, both the CCW and CW residence times increase, while the global distribution of motor bias is less affected. Finally, the fusions to MotB induce a novel BFM dynamic that involves an asymmetry in the speeds attained in opposing rotational directions, depending on the motor’s bias. These effects of the fluorescent fusion proteins on the BFM could be partially restored by a rigid linker (EAAAK)3 inserted at the fusion point. Together, these findings reveal how a fluorescent protein fusion that does not abolish function completely can modulate biologically relevant dynamics of a protein complex and even induce new behaviour, all of which can be partially relieved by the incorporation of a linker.

The fact that an FP fused to the stator causes an overall decrease of speed could be explained by a variety of mechanisms. For example, the tags could perturb ion translocation, hinder the stator conformational changes involved in force generation, inhibit the interaction between the stator and rotor, introduce an extra drag in the rotor, or interfere with stator recruitment. While our measurements cannot discriminate a single mechanism, our analysis shows that the number of stators is not affected in tagged motors, ruling out an interference with stator recruitment.

The fusion proteins also lowered the switching frequency. It has previously been observed that the switching frequency is dependent upon motor torque and rotation speed [49], and it has been proposed that the conformation of the switch complex is dependent upon the interactions with the stators in a torque-dependent manner [50,51]. It is thus plausible that the observed decrease in switch frequency is due to a decrease in the single stator torque due to the FP fusion. Alternatively, it could be that the fusion protein causes a reduced switching frequency by direct interactions with the governing structural elements.

The unexpected discovery that FP-MotB fusions cause a bias-dependent speed asymmetry indicates that the fluorescent protein interacts with a yet unspecified asymmetry in the stator-rotor interaction that may or may not be present in the native BFM. Importantly, we note that the mechanisms suggested above affect speed symmetrically in both directions, and cannot account alone for the observations of speed asymmetry. Further study is needed to resolve the exact mechanism that gives rise to the observed bias-dependent speed asymmetry. In SI 4.4 we speculate about possible mechanisms that could be responsible for this asymmetry.

It is increasingly clear that the selection of a proper linker is an important element of FPF design [10,11] While properties such as linker flexibility, length, and hydrophobicity are important, predicting the success of a linker *a priori* is difficult. Here, a longer linker performed better than a shorter linker, and a rigid linker performed better than a flexible linker. We hypothesize that these particular properties allow for a sufficient spatial separation between MotB and the FP, thereby reducing the interactions between the FP and the force-generating MotB/FliG interface.

In science, the act of observing often impinges upon the phenomenon of interest. The power of fluorescent protein fusions as a tool for the study of protein function hinges on our understanding of their side effects. Here we have thoroughly characterized these effects for the stator of the BFM. The findings advance our ability to study protein-complex dynamics while imparting minimal perturbation.

## 4 Methods and Materials

### 4.1 Bacteria and culture preparation

All the *E. coli* strains used are detailed in Table 1. Three FPs (eGFP, YPet, and Dendra2) were fused either directly to the N-terminus of MotB or with a linker in between, as detailed below. A cassette containing the *MotA* and FP-fused *MotB* genes were cloned downstream of the promotor in plasmid pBAD33 (ara, araC, pACYC184/p15A) [52] and transformed into the cells. Cells were grown at 33°C in tryptone broth with chloramphenicol (34 ug/ml) and either 0.13mM or 6.5mM of L-arabinose to an OD_600_ of 0.55-0.65.

For all of the individual cell measurements, the flagella were mechanically sheared [56] before the cells were centrifuged and resuspended in motility buffer (10 mM potassium phosphate, 0.1 mM EDTA, 10 mM lactic acid, pH 7.0). Measurements were carried out at 20°C in a simple flow chamber made by two coverslips separated by a layer of parafilm.

The population-level motility was measured by observing the chemotactic population front on soft (0.25%) agar plates. As cells metabolize nutrients in the agar, they create an attractant gradient and, if chemotactic, swim out from the point of inoculation, forming an expanding ring whose diameter was measured after 8h (each measurement was repeated in triplicate, see Fig 1B). In order to choose the best linker, the population motility was measured for Dendra2 stators joined to four different linkers of the following sequences: (GGGGS), (EAAAK), (GGGGS)_3_, and (EAAAK)_3_.

### 4.2 Motor rotation measurements

Speed measurements of individual motors were performed using a tethered-bead assay [56], as follows. Bacterial cells were immobilized to poly-L-Lysine (Sigma) coated coverslips, and polystyrene beads (1.1 *μ*m diameter, Sigma) were allowed to spontaneously attach to truncated ‘sticky’ (FliC^st^) filaments. Motor rotation was measured by tracking the rotation of the bead, which was monitored with a bright-field laser microscope setup [57]. Images of the bead hologram were recorded by a CMOS camera (Optronics) at the rate of 300 or 500 frames per second for 3 to 4 minutes. Where tested, the linker joining MotB to the FP was (EAAAK)_3_.

Custom Labview and Python software was used to track the bead with nm resolution, correct for sample drift, and fit an ellipse to the trajectory (considering the ellipse as the projection of a tilted circle). The angle of the bead with respect to the ellipse determined the angular position, and the angular speed of motor rotation was calculated from the time derivative of the angular position. The speed was then median-filtered with a window of 70 ms. Positive (negative) speed indicates CCW (CW) rotation. From the speed trace, motor bias was measured as the proportion of time spent rotating CCW. A motor that spent more than 50% of the time rotating in the CCW (CW) direction is referred to as CCW- (CW-) biased. Speed histograms of individual motors were constructed with a sub-Hz bin-width. The individual normalized histograms are shown as light lines in Fig 2 as probability densities (in logarithmic scale). The global average distributions are shown by the thick lines. The line color of the speed distributions in Fig 2 reflects the motor bias, blue and red indicating CCW-and CW-biased motors, respectively. Rotational switching events were detected from the median-filtered speed trace, based on an algorithm which finds the crossing of two thresholds, set at 2/3 of the mean speed in each direction [58]. The switching frequency of a motor was calculated by the number of detected switching events divided by the duration of the measurement.

### 4.3 Estimate of single stator torque contribution

The torque generated by the motor was calculated as the product of the drag coefficient of the bead [59] and the speed. In several motors at steady-state (here ΔCheY to avoid complications due to switching) we could observe jumps between discrete torque levels (one example is shown in Fig 4A). For each of these traces, the histogram of motor torque had multiple peaks, which we fit with a multiple Gaussian fit. Under the common assumption that discrete changes in motor torque are due to a change in stator number [21, 37, 44, 56, 60, 61], expected due to stator-turnover [15], the distance in the histogram from one peak to the next represents the torque contributed by a single stator. The distance between neighboring Gaussians was calculated for each individual trace, and the distribution of such single-stator torque contributions for all the measured motors for WT and all of the FP-MotB strains is shown in Fig 4B. Finally, a Gaussian was fit to this distribution to determine the average single-stator torque contribution.

### 4.4 Fluorescence microscopy measurements

Fluorescence measurements of individual motors were performed using a tethered-cell assay [62], as follows. Bacterial cells were allowed to spontaneously adhere to the glass coverslip via the truncated hydrophobic filament. Cells which tethered by a single filament rotated around the axis of the corresponding flagellar motor (Fig 1C).

Fluorescence excitation was performed at the wavelength of 488 nm in a TIRF configuration (at a power of ~200 W/cm^2^, measured before the objective) and emission was detected by an EMCCD camera (Andor) in the range of 500-550 nm. The emission of Dendra2, a photo-switchable fluorophore, was detected in its unconverted form. Tethered cells with labeled motors were often observed rotating around a bright fluorescent spot, indicating the location of the functional motor. The fluorescence intensity of nine pixels around the center of rotation was summed to obtain the motor signal *S_m_*. Only fluorescent spots which coincided with the center of rotation of a rotating cell were analyzed, to avoid false positives due to e.g. possible clusters of FPs. Due to rotation and blur, the auto-fluorescence of a rotating cell was not possible to quantify reliably; therefore, the auto-fluorescence of ~ 20 stuck cells in the same field of view was averaged to obtain the noise level *S_n_*. Fig 1D shows the signal-to-noise ratio, defined by SNR = *S_m_*/*S_n_*, for the different strains. An SNR of one indicates that the motor is no more fluorescent than the autofluorescent cellular background. An SNR greater than one indicates the presence of fluorophores at the motor, indicating proper folding of the fluorescent protein and successful integration of the stator to the motor.

## Supporting information

**S1 Fig. Fluorescence time traces.**

**S2 Fig. Speed histograms for the two induction levels used.**

**S3 Fig. Switching at the two induction levels tested.**

**S4 Fig. Residence time distributions at the two induction levels tested.**

**S5 Fig. Chemotaxis motility comparisons of the Dendra2 fusion stators.**

**S6 Fig. The (EAAAK)_3_ linker improves the performance of YPet and eGFP motors.**

**S1 Appendix. Possible origins of the speed asymmetry in switching.** One first hypothetical mechanism that could explain the asymmetry in switching can be an asymmetric effect of the tags on the power stroke of the stator. Following a recent model proposing that the two pairs of the cytoplasmic loops of MotA may drive the rotation in opposite directions [63], one could speculate that the presence of two tags on MotB could perturb one pair of the cytoplasmic MotA loops more severely than the other, decreasing the efficiency of torque generation in one direction more than in the other. In this scenario, different stators should then assemble with the same orientation of their efficient MotA pairs with respect to FliG, otherwise a random proportion of efficient and inefficient MotA pairs of different stators aligned with either FliG orientations (CW or CCW) would not give the asymmetrical switch which we systematically observe.

A second hypothetical mechanism could be to assume that the presence of the tags, in proximity of the C ring, prevents the complete conformational switch of the nearest FliG unit. Here the tag, blocking FliG to complete its transition, would reduce, but not necessarily eliminate, the torque in the corresponding conformation. This mechanism relies on the hypothesis of the existence of multiple intermediate functional conformations of FliG.

A third hypothetical possibility could come from the observation that the switching frequency of tagged motors decreases, while the bias is less affected, indicating that the energy barrier between the two conformational states of the FliG protein of the C-ring is increased by the presence of the tag. This could have the effect, in average, to lock a few FliG units in their conformations, depending on the motor bias. The observations could be explained in terms of the conformational spread model, in which it has been shown that locking only a few FliG units can have a strong influence in the switching dynamics of the motor, in terms of bias, maximum speed and residence times [64]. The above speculations remain to be tested.

## Acknowledgments

We thank RM Berry for the gift of several strains, and G Labesse for helpful discussions. The CBS is a member of the France-BioImaging (FBI) and the French Infrastructure for Integrated Structural Biology (FRISBI), 2 national infrastructures supported by the French National Research Agency (ANR-10-INBS-04-01 and ANR-10-INBS-05, respectively). We acknowledge funding from the European Research Council under the European Union’s Seventh Framework Programme (FP/2007-2013)/ERC Grant Agreement n. 306475.

---

**Fig S1.**
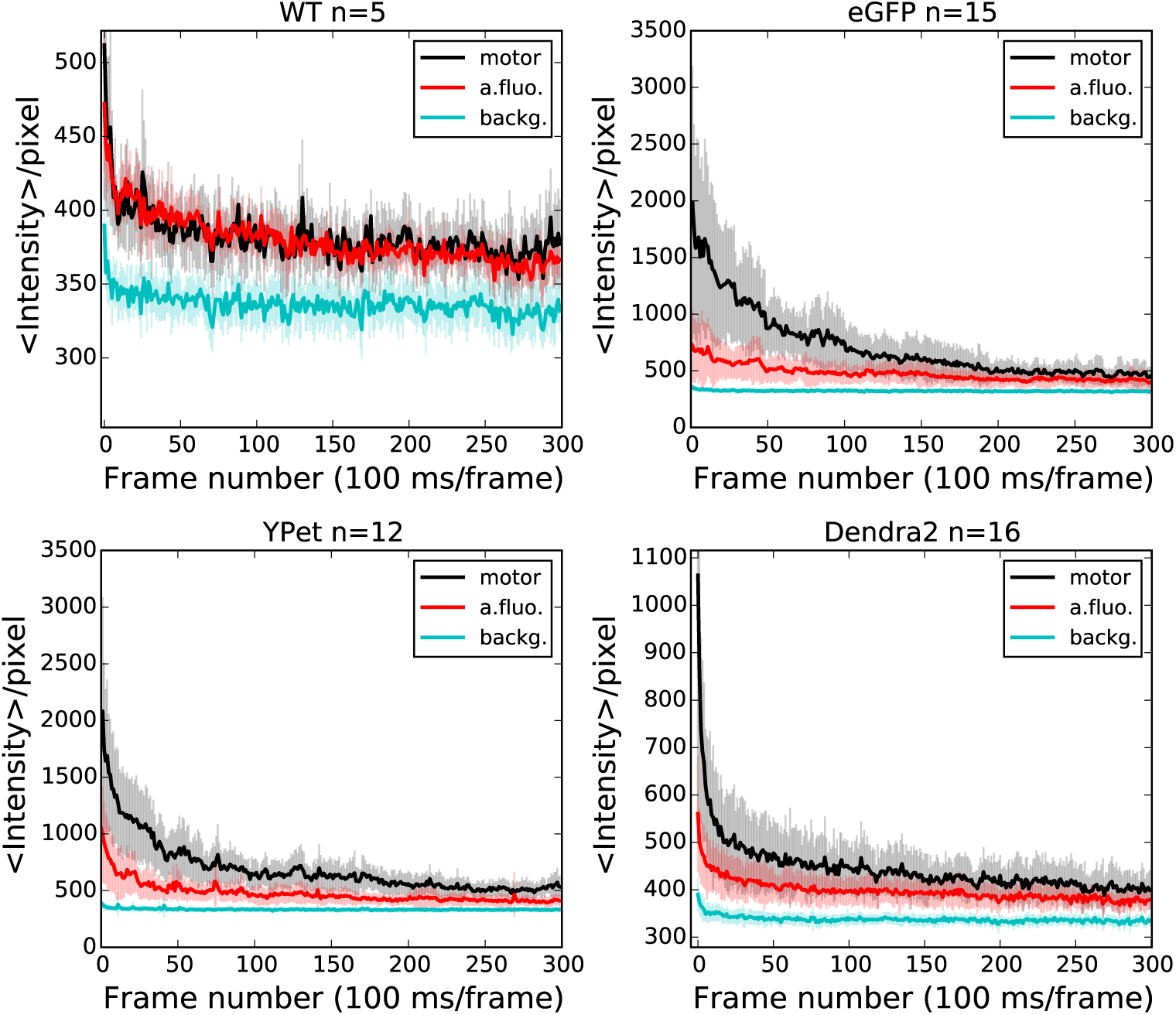
Fluorescence time traces. For each strain we have measured the time evolution of the fluorescence signal from a 3x3 pixels region around an active motor (black line) on *n* rotating tethered cells (the strain and *n* are indicated in the title of each panel). The cellular auto-fluorescence signal (red) is measured on 3x3 pixel regions selected over several (~ 20) stuck cells in the same field of view. The background signal (cyan) is measured on regions not occupied by cells. [Arabinose] = 0.13 mM. Error bars indicate the standard deviation.

**Fig S2.**
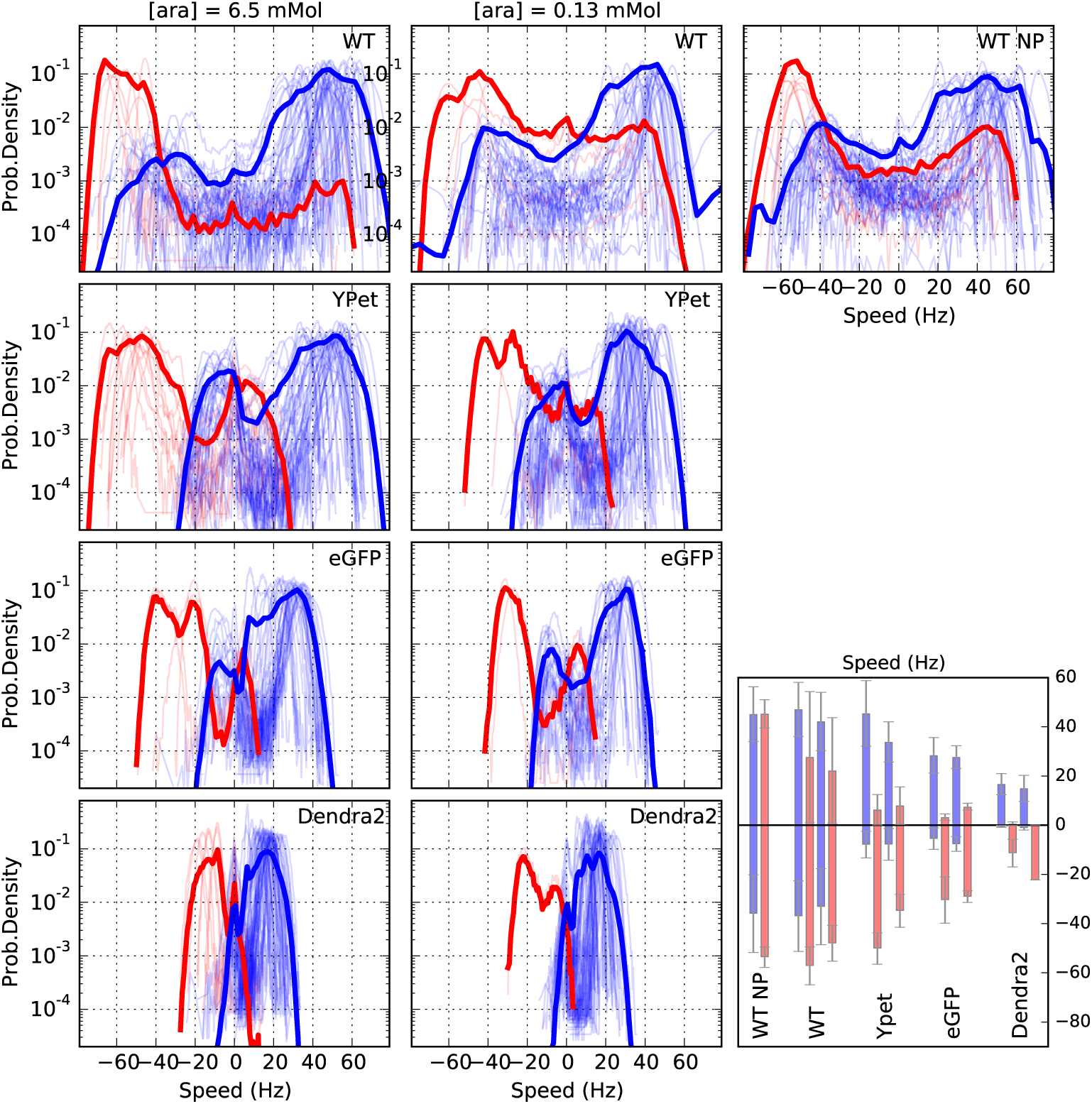
Speed histograms for the two induction levels used (first column: [Ara] = 6.5 mM, second column: [Ara] = 0.13 mM, while WTNP corresponds to the native expression). As in Fig.2 of the main text, positive (negative) speed indicates CCW (CW) direction, and blue (red) indicates CCW (CW) biased motors. Individual thin lines show individual motor measurements, and the thick lines show the average of all measurements. The bar plot at the bottom right summarizes the mean speed and standard deviation for each strain for both expression levels, for CCW and CW biased motors (blue and red bars, respectively). For the strains with the stators on a plasmid, WT, YPet, eGFP, and Dendra2, the first two bars correspond to high induction (6.5 mM), the second two bars to low induction (0.13 mM).

**Fig S3.**
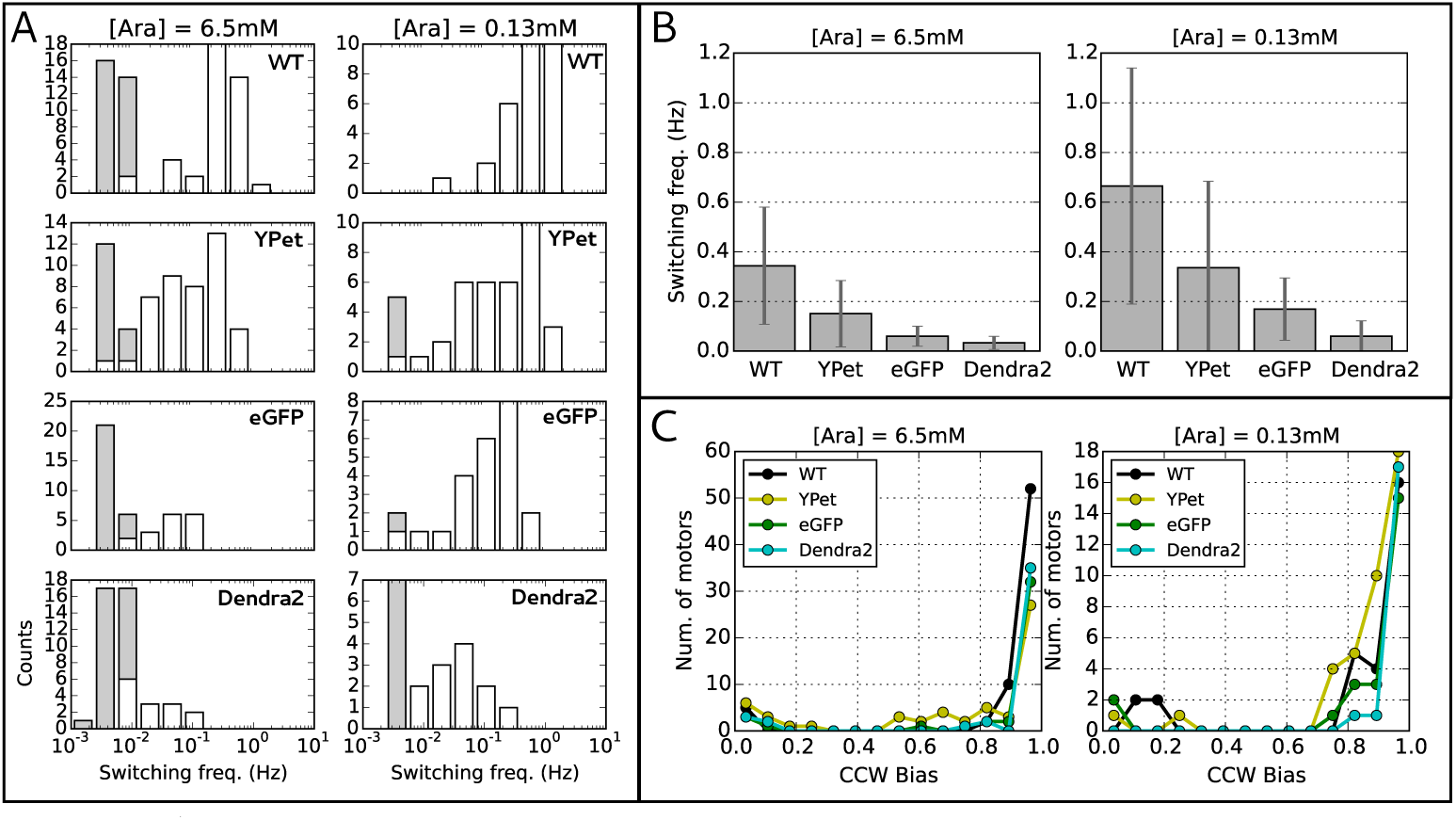
A) Distributions of switching frequency for all the strains, at the two induction levels tested (indicated in the title of the two columns). As in Fig 3, the gray bars indicate motors which did not switch during the measurement, so their switching frequency has been set equal to twice the inverse of the measurement time, and it should be considered as an upper limit. B) Summary of switching frequency mean and standard deviation for all the strains at the two induction levels tested. C) Distributions of the CCW bias for all the strains at the two induction levels.

**Fig S4.**
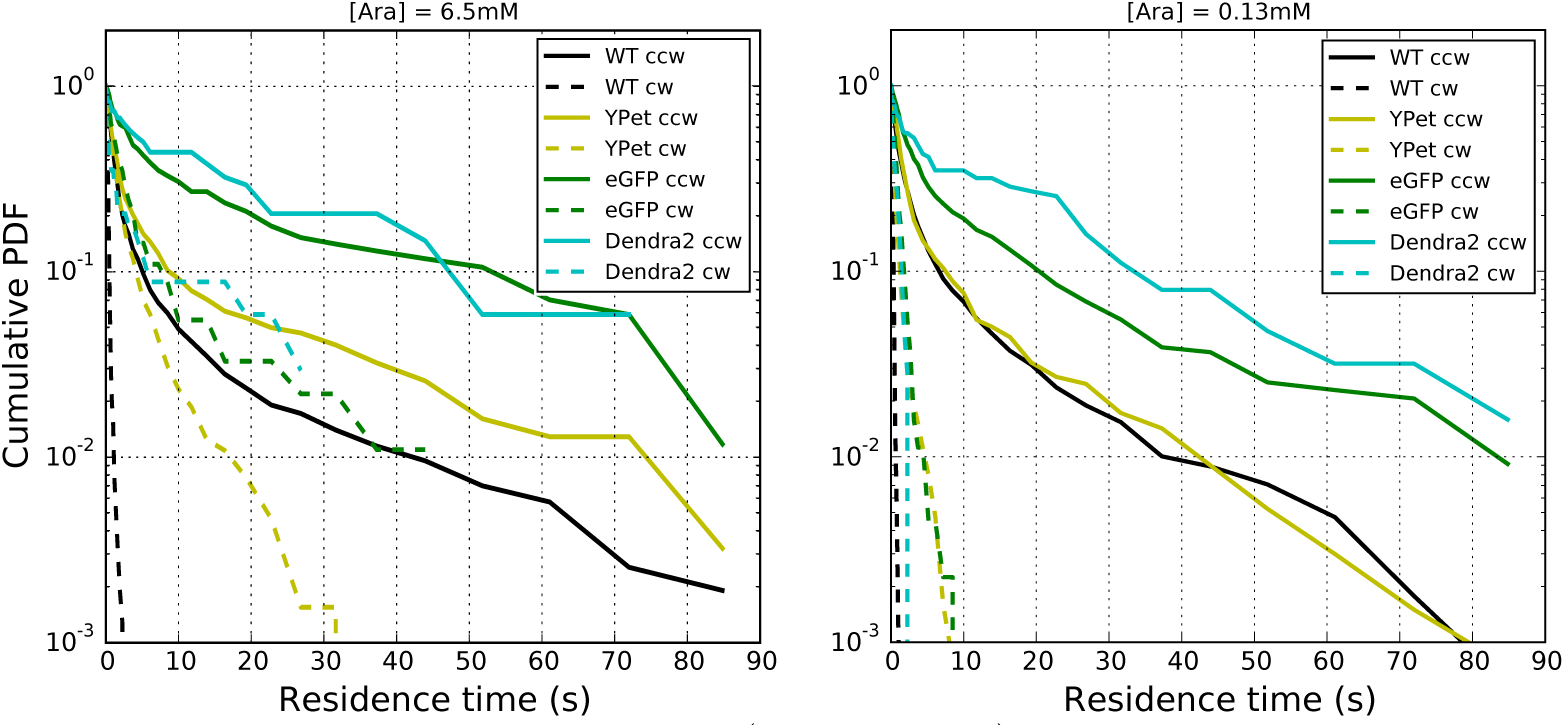
Inverse cumulative distribution (as in Fig. 3B) of the residence times for all CCW biased motors at the two induction levels tested, indicated in the figure titles.

**Fig S5.**
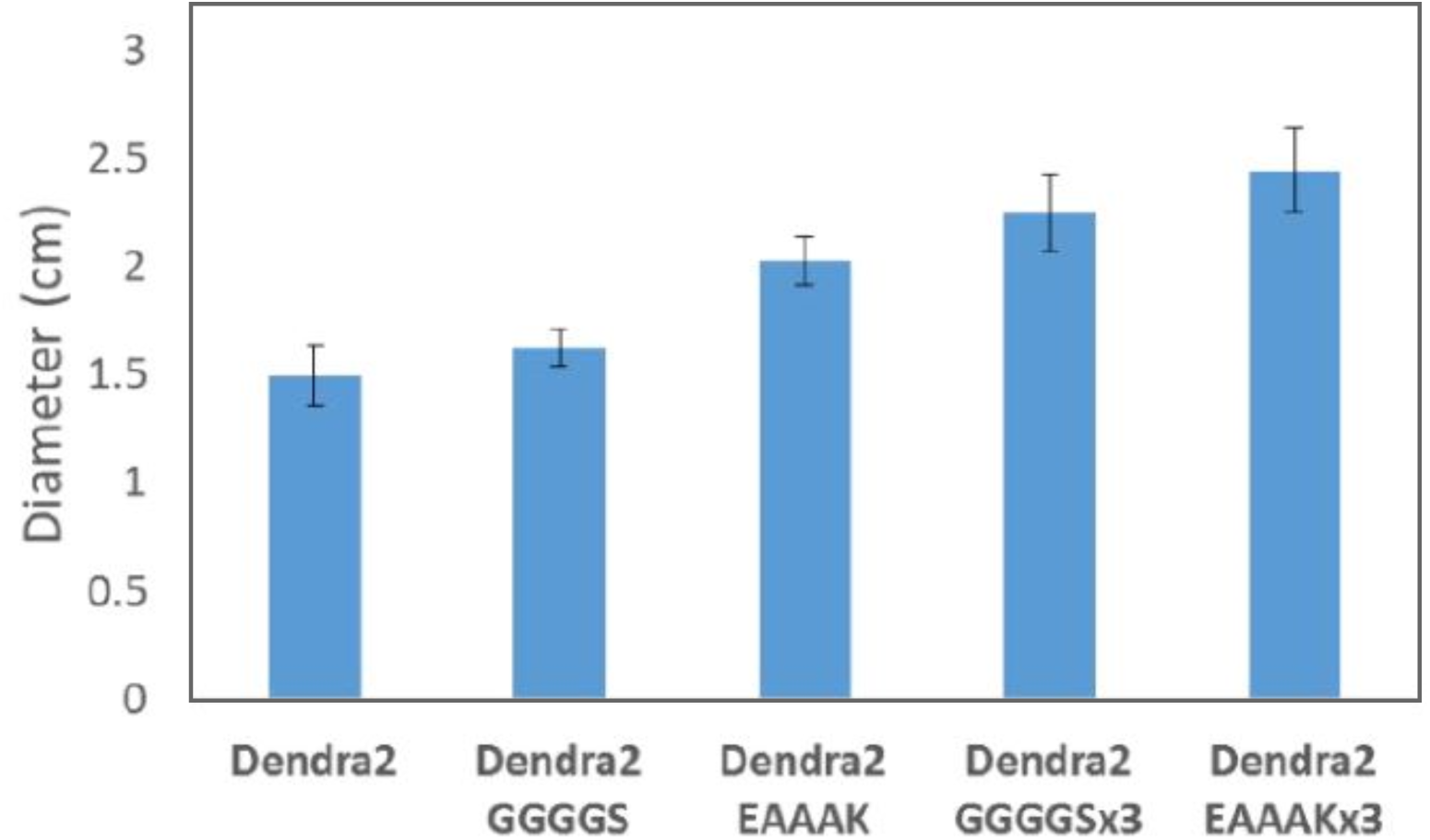
Chemotaxis motility comparisons of the Dendra2 fusion stators. Error bars give standard deviation over three measurements. The rigid linker (EAAAK) yields a larger improvement than the flexible (GGGGS) linker, and the triple repeat yields a larger improvement than the single.

**Fig S6.**
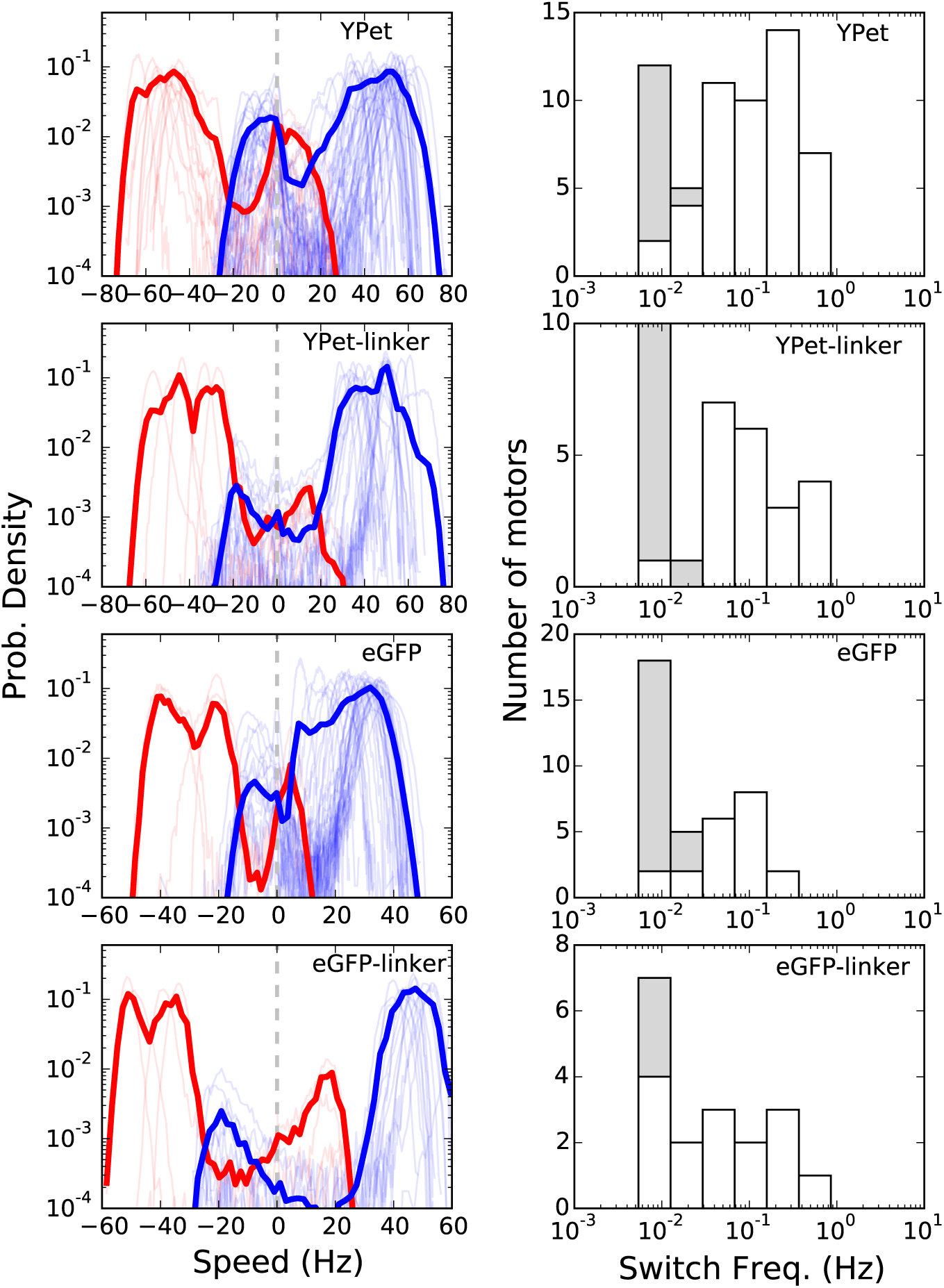
The (EAAAK)_3_ linker improves the performance of YPet and eGFP motors. Left column: Distributions of the speed measured in YPet and eGFP motors, in the presence and absence of the linker, as indicated in each panel (the data in the absence of linkers are the same as in Fig. 2, shown as a reference). Right column: distributions of the switching frequency measured for the same strains. As in Fig. 3 and Fig. S3, the gray bars indicate motors which did not switch during the measurement, so their switching frequency should be considered as an upper limit. [Ara] = 6.5 mM

